# ΔFBA: Predicting metabolic flux alterations using genome-scale metabolic models and differential transcriptomic data

**DOI:** 10.1101/2021.01.18.427188

**Authors:** Sudharshan Ravi, Rudiyanto Gunawan

## Abstract

Genome-scale metabolic models (GEMs) provide a powerful framework for simulating the entire set of biochemical reactions occurring in a cell. Constraint-based modeling tools like flux balance analysis (FBA) developed for the purposes of predicting metabolic flux distribution using GEMs face considerable difficulties in estimating metabolic flux alterations between experimental conditions. Particularly, the most appropriate metabolic objective for FBA is not always obvious, likely context-specific, and not necessarily the same between conditions. Here, we propose a new method, called ΔFBA (deltaFBA), that employs constraint-based modeling, in combination with differential gene expression data, to evaluate changes in the intracellular flux distribution between two conditions. Notably, ΔFBA does not require specifying the cellular objective to produce the flux change predictions. We showcased the performance of ΔFBA through several case studies involving the prediction of metabolic alterations caused by genetic and environmental perturbations in *Escherichia coli* and caused by Type-2 diabetes in human muscle.

## Introduction

In the post-genomic era, there has been intense efforts directed toward the reconstruction of genome-scale models of cellular networks. An important portion of these efforts focuses on metabolic networks due to the significance of cellular metabolism for understanding diseases such as cancer ^1–4^ as well as for metabolic engineering applications in biomanufacturing ^5^. Recent advances in high-throughput sequencing technologies, gene functional annotation, and metabolic pathway databases, and developments of algorithms for mapping gene-protein-reaction (GPR) associations and identifying missing metabolic reactions systematically (gap-filling), have enabled the reconstruction of thousands of genome-scale metabolic models (GEMs), from single cell organisms to human ^6,7^. A GEM provides the gene-protein-reaction associations that encompass the set of metabolites and metabolic reactions in an organism as prescribed by its genome. Concurrent with these developments are the creation of efficacious algorithms that use GEMs to predict intracellular metabolic fluxes – i.e. the rates of metabolic reactions – and how these fluxes vary under different environmental, genetic, and disease conditions ^8–10^.

A prominent class of algorithms based on a constrained-based modeling technique, called flux balance analysis (FBA), have flourished due to its ease of formulation and flexibility, using the stoichiometric coefficients of the metabolic reactions in a GEM, an assumed cellular objective such as maximization of biomass production, and various experimental data on metabolic capabilities and constraints of the cells, to predict metabolic fluxes ^11^. Although FBA is effective in handling large networks and predicting cell behavior in many metabolic engineering studies ^12–15^, considerable uncertainty still remains about the appropriate choice of cellular objective for different conditions and cell types – a choice that requires expert knowledge of the cells and their phenotype in a given condition. Such an issue is particularly prominent for complex organisms such as human. Moreover, the flux solutions produced by applying FBA have degeneracy – that is, multiple solutions exist that give the same cellular objective value ^16^. Not to mention, standard FBA often produces biologically unrealistic flux solutions ^17,18^.

Driven by the increasing ease and availability of whole-genome omics profiling data, a multitude of FBA-based algorithms have been developed to incorporate additional datasets to create context-specific metabolic networks and to improve flux prediction accuracy ^19–26^. Several of these methods, such as GIMME (Gene Inactivity Moderated by Metabolism and Expression) ^19^, iMAT (integrative Metabolic Analysis Tools) ^21^, and MADE (Metabolic Adjustment by Differential Expression) ^25^, are based on maximizing the consistency between the predicted flux distribution and the mRNA transcript abundance of metabolic genes, where the higher the transcript level of an enzyme, the larger should the flux of the corresponding reactions. Other methods, such as E-Flux ^23^, use data of transcript abundance for setting the bounds on reaction fluxes. More recent methods, such as ME-model ^27^ and GECKO ^28^, combine FBA with an explicit modeling of enzyme / protein expression and thus, are able to directly account for protein abundance. Thermodynamics constraints have also been integrated with the FBA to eliminate thermodynamically infeasible fluxes, and at the same time enable the integration of metabolite concentration data, as done in recent methods such as ETFL ^29^.

A number of methods focus on using differential expression data between two conditions (e.g., perturbation vs. control) to predict metabolic alterations – a particular focus of our study. The method Relative Expression and Metabolic Integration (REMI) ^30^ used differential expression of transcriptome and metabolome to estimate metabolic flux profiles in *Escherichia coli* under varying dilution and genetic perturbations. The method relies on maximizing the agreement between the fold-changes of metabolic fluxes and the fold-changes of enzyme expressions between two conditions. The metabolome data, if available, are used to determine the flux directionality using reaction thermodynamics. Among the alternative flux solutions, the L1-norm minimal solution is adopted to give a representative flux distribution. Another method by Zhu *et al.* ^31^ employed a softer definition when assessing consistency between the metabolic fluxes and enzyme differential expressions, where only the sign of the differences needs to agree. The method provides a qualitative determination of metabolic flux changes by determining the maximum and minimum flux through each reaction in the GEM. Both of the above methods generate metabolic flux predictions for each condition in comparison, and like standard FBA, both methods require an assumption on the cell’s metabolic objective. Generally, model prediction inaccuracy is amplified when evaluating the differences between two model predictions. Another related method MOOMIN ^32^ uses a Bayesian approach to integrate differential gene expression profiles with GEMs to predict the qualitative change in the metabolic fluxes—increased, decreased or no change.

In this work, we developed ΔFBA (deltaFBA) for predicting the metabolic flux difference given a GEM and differential transcriptomic data between two conditions. ΔFBA relies on a constrained-based model that governs the balance of flux difference in the GEM, while maximizing the consistency and minimizing inconsistency between the flux alterations and the gene expression changes. ΔFBA is created in MATLAB to work with the COnstraint-Based Reconstruction and Analysis (COBRA) toolbox ^33^. We applied the ΔFBA to analyze the metabolic changes of *Escherichia coli* in response to environmental and genetic perturbations using data from the studies of Ishii *et al.* ^34^ and Gerosa *et al.* ^35^. We compared the performance of ΔFBA to REMI for evaluating flux differences between conditions. We also demonstrated the application of ΔFBA to human GEM, specifically evaluating the metabolic alterations associated with type-2-diabetes in skeletal muscle using myocyte-specific GEM ^36^.

## Materials and Methods

### Method Formulation

ΔFBA generates a prediction for metabolic flux changes between a pair of conditions, such as treated vs. untreated or mutant vs. wild-type strains. In the following, we use the superscript *C* to denote the control (reference) condition and P to denote the perturbed condition. ΔFBA enforces that the flux changes Δ***ν* = (*ν*^P^ − *ν*^C^)** for each metabolite are balanced, as follows:

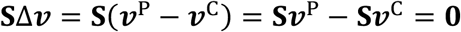

where **S** denotes the *m* **×***n* stoichiometric matrix for *m* metabolites that are involved in *n* metabolic reactions in the GEM, and ***ν*** and Δ***ν*** denote the vector of metabolic fluxes and flux changes, respectively. The flux balance above is a direct consequence of the steady-state flux balance in each condition, i.e. **S = 0**

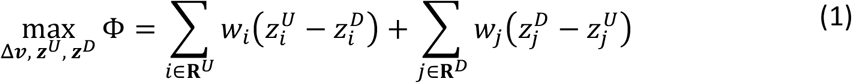

subject to:

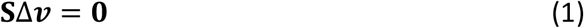

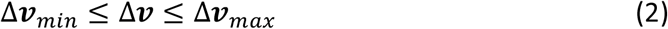

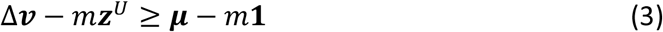

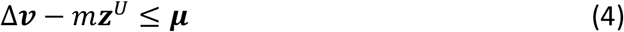

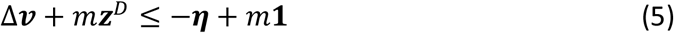

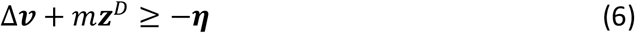

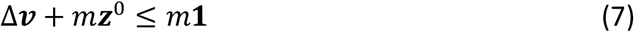

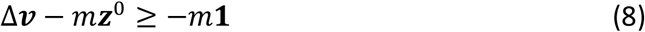

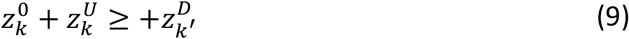

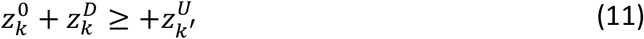

The weighting coefficients ***wi*** allows users to prioritize certain reactions (e.g., reaction(s) corresponding to gene knock-out(s)) for (in)consistency. The set of upregulated reactions **R*^U^*** and the set of downregulated reactions **R*^D^*** are user-defined inputs. Typically, the sets **R*^U^*** and **R*^D^*** include reactions with statistically significant increase and decrease in gene expression between the perturbed condition and the control, respectively. Equation (2) specifies the upper and lower bounds for the flux change, which can be set based on experimental data (e.g. the change in the biomass production or growth rates The weighting coefficients, or based on the corresponding bounds from the perturbed **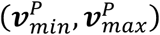** and the control condition **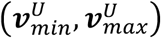** as follows:

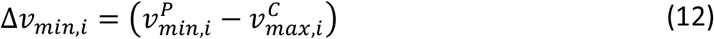

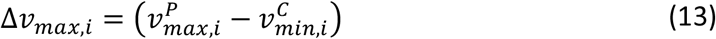

The MILP above will generate flux change prediction Δ***ν*** whose signs (positive and negative) agree with the direction of the gene expression changes. Specifically, the binary decision variables **z*^U^*, z*^D^***, and **z^0^** in the MILP formulation set the sign (directionality) of the flux changes Δ***ν***. When **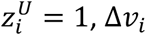** takes a positive value above a threshold ***μ*(**, as specified by Equations (3)-(4). Vice versa, when **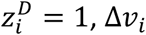** takes a negative value below a threshold ***η*(**, as specified by Equations (5)-(6). The thresholds ***μ*_*i*_** and ***η*_*i*_** for the positive and negative flux changes, respectively, are user-defined parameters. For example, the thresholds can be set to the same constant value ***ε***, or to a value that scales with the fold-change reaction expression **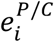** (see Supplementary Materials), as follows:

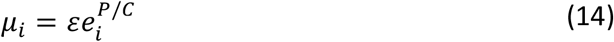

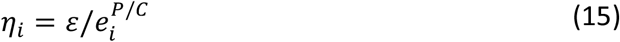

Clearly, **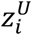** and **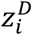** cannot simultaneously be equal to 1 since no feasible Δ***ν*(** exists that simultaneously satisfy Equations (3)-(6). The decision variable ***z*^0^** is used to force select reactions to have zero flux change value, as specified by Equations (7)-(8). Together with Equation (9)-(11), the forward and reverse halves of a reversible reaction, Δ***ν*3** and Δ***ν*3#**, are prevented to simultaneously have non-zero values, so as to reduce degeneracy of the flux change solution Δ***ν***.

Given the degrees of freedom in GEMs for Δ***ν***, many equivalent optimal solutions often exist that give the same objective function value Φ^*^ as specified in Equation **Error! Reference source not found.**. By assuming parsimony in Δ***ν*** – that is, Δ***ν*** is minimal between the perturbed and control condition, a two-step optimization procedure is implemented in ΔFBA. The first step is to maximize consistency with gene expression changes as prescribed in Equations **Error! Reference source not found.**-(11) with the maximum objective function value denoted by Φ^*^. The second step is to produce an L2 norm minimal solution for Δ***ν***, as follows:

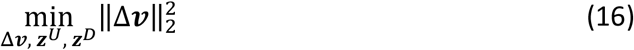

subject to the same constraints in Equations (1)-(11) while achieving the same level of consistency Φ^*^, implemented by the additional constraint:

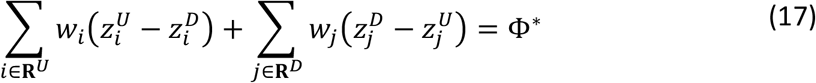

The L2 minimization is based on the premise that the perturbed metabolic fluxes should not deviate far from the control condition, similar to the method called Minimization of Metabolic Adjustment (MOMA) ^37^. An alternative to L2-norm minimization is L1-norm minimization, which is analogous to maximizing sparsity of Δ***ν***. The L1-norm minimization was previously used in the parsimonious FBA (pFBA) method, but such an approach often still leads to multiple degenerate solutions. On the other hand, the L2-norm minimization will produce a unique solution, but the mixed integer quadratic optimization that is required to find the solution may have high computational requirement.

ΔFBA is available as MATLAB scripts and are compatible with the COBRA toolbox. ΔFBA requires Gurobi optimizer (http://www.gurobi.com) as a pre-requisite. ΔFBA has been tested on a Windows PC using a 6-core Intel Xeon (2146G) Processor with 16 GB RAM.

### Case studies: Data and Implementation

The first case study involved the response of *E.* coli’s metabolism to genetic (single-gene deletions) and environmental perturbations (dilution rates) performed by Ishii *et al*. ^34^. The study provided ^13^C-based flux data and RT-PCR mRNA abundances for the central carbon metabolism, pentose phosphate pathway (PPP), and the tricarboxylic acid (TCA) cycle for wild-type K12 *E. coli* culture in chemostat under different dilution rates (0.1, 0.2, 0.4, 0.5, and 0.7 hours^−1^) and for 24 single-gene perturbations along the glycolysis and PPP ^34^. The global transcriptional response was only captured for 5 of the 24 single-gene deletions (*pgm, pgi, gapC, zwf* and *rpe*) and two of the 4 dilution conditions (0.5 and 0.7 hours^−1^). The differential (fold-change) gene expression levels were computed with respect to the control condition, set to be wild-type K12 *E. coli* cultured at a dilution rate of 0.2 h^-1^. The differential (fold-change) reaction expressions were subsequently evaluated based on the fold-change gene expression using the GPR Max/Min rule available in the COBRA toolbox (MATLAB) ^38^.

For samples with only RT-PCR mRNA abundance data, the set of up- and downregulated reactions, **R*^U^*** and **R*^D^***, included all reactions with fold-change reaction expressions higher than 1 and those with fold-change lower than 1, respectively. In the additional analyses for samples with whole-genome transcriptome data, the set of up- and downregulated reactions, **R*^U^*** and **R*^D^***, respectively, were taken from the top and bottom 5^th^ percentile of the differential reaction expressions. The differences of the measured cell specific glucose uptake rates between perturbed and control experiments were used as constraints. ΔFBA was applied using the two-step optimization with the L2 norm minimization, as described above.

The second case study came from a study of *E. coli* growth on 8 different carbon sources performed by Gerosa *et al*. ^35^. Unprocessed global transcriptomic data were obtained from ArrayExpress (E-MTAB-3392) ^39^, and differential expression analyses between every pair of carbon sources were evaluated using the *Limma* package in R ^40^. As before, the fold-change reaction expressions were computed based on fold-change in the global gene expression using the Max-Min GPR rule using COBRA toolbox ^38^. The up- and downregulated set of reactions were taken from the top and bottom 5^th^ percentile of the differential reaction expressions. In addition, cell culture data on specific growth rates were used to compute the flux change bounds for biomass production rate. The uptake rates of the carbon source changes were also incorporated as constraints. We implemented the two-step optimization of ΔFBA using L2 norm minimization.

The third case study came from two studies of skeletal muscle tissue metabolism in type-2 diabetes (T2D) patients by van Tienen *et al.* ^41^ and Jin *et al.* ^42^. The microarray gene expression datasets were obtained from GEO (GSE19420 ^41^ and GSE25462 ^42,43^) and the differential (fold-change) expression of genes for each dataset were computed using the *Limma* package in R ^40^. The fold-change reaction expressions were computed based on the differential gene expression using the Max/Min GPR rule. In the absence of additional constraints in the form of exchange fluxes or growth characteristics, we set the up- and downregulated reactions from the top and bottom 25^th^-percentile in differential reaction expressions, rather than the 5^th^ percentile threshold used in *E. coli* case studies above, so as to incorporate more differentially expressed transcripts. We implemented an L1-norm minimization in the second step of ΔFBA to reduce computational complexity (time) due to the large number of constraints associated with the differential reaction expressions.

### Implementation of REMI

The method Relative Expression and Metabolomic Integrations (REMI) was developed for predicting individual flux distributions of a pair of conditions (***ν*^P^** and ***ν*^C^**) using multi-omics dataset. The toolbox was downloaded from https://github.com/EP-LCSB/remi. The differential gene expressions in each case study were obtained as described above. The mapping from differential gene expression to the corresponding reaction expressions were done using the procedure detailed in REMI ^30^. Briefly, the authors followed the implementation of Fang *et al.* ^33^ to translate gene expression ratios to obtain reaction expression ratios. When several enzyme subunits are required for a reaction, a geometric mean of expression ratios is chosen to represent the reaction ratio. In the case where multiple isozymes catalyze a reaction, the arithmetic mean of the individual expression ratios of the isozymes is used for the reaction ratio. The set of up- and down-regulated reactions **R***^U^* and **R*^D^*** were taken from the computed differential reaction expressions as in ΔFBA implementation. Unlike ΔFBA, REMI produces solutions for the metabolic fluxes of perturbed ***ν*^P^** and control condition ***ν*^C^**. For comparison, we evaluated the flux change predicted by REMI by taking the difference: Δ***ν* = *ν*^P^ − *ν*^C^**.

### Performance evaluation

The quantitative agreement between the measured flux difference Δ***ν*^9^** and the predicted flux changes Δ***ν*^*^** was assessed by using two accuracy metrics: uncentered Pearson correlation coefficient and normalized root mean square error (NRMSE). The uncentered Pearson correlation coefficient ***ρ*** was computed as follows

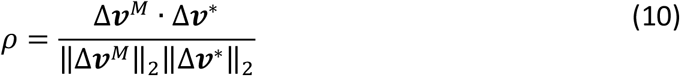

Meanwhile, the NRMSE was according to the following equation – using *tdStats* package in R:

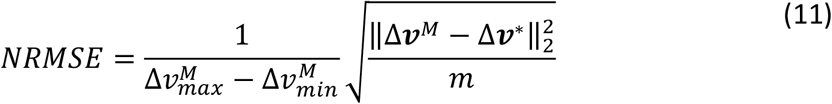

Besides the quantitative agreement in flux changes, we also evaluated the qualitative agreement by comparing the signs of the flux changes between experimental measurements and predictions. To this end, we discretized the measured and predicted flux changes into +1, 0, and −1, to describe upregulated, no change, and downregulated reactions, respectively. The agreement in the direction of the flux changes was evaluated as the number of correct sign predictions divided by the total number of fluxes.

### Metabolic subsystem enrichment analysis

The flux differences obtained from applying ΔFBA were first filtered according to the directionality of their change. The significantly altered fluxes (**|**Δ***νi*| > *ε***) were grouped based on the subsystem to which the fluxes belong. A Fisher exact test (*fisher.test* function in the R-package) was used in determining over-represented subsystems in upregulated (positive change) and downregulated (negative change) fluxes. The statistical significance *p*-values were corrected for multiple hypothesis testing using the *p.adjust* function in R.

## Results

### Escherichia coli response to genetic and environmental variations

Ishii *et al.* ^34^ studied the robustness of *E. coli* K12 metabolism in chemostat in response to changes in dilution rates and to gene deletions. The study generated multi-omics data, including transcriptomic, proteomic, metabolomic, and ^13^C metabolic fluxes, and demonstrated the remarkable ability *of E.* coli to reroute its metabolic fluxes to maintain metabolic homeostasis in response to environmental and genetic perturbations. Only a small fraction of variation in the measured flux ratios can be explained by the fold-change in reaction expressions, as indicated by the low coefficient of determinations R^2^, regardless of the GPR mapping procedures in ΔFBA and REMI (see **Figure 1A**). The weak agreement between reaction expressions and metabolic fluxes motivates using a system-oriented approach that consider the global network changes based on GEMs ^44^. We applied ΔFBA using *E. coli*’s *iJO1366* GEM to predict the metabolic flux shifts from the control condition (wild-type K12 at 0.2 hour^-1^ dilution rate), caused by alterations in dilution rates (0.1, 0.4, 0.5, and 0.7 hours^−1^) and by 24 single-gene deletions (*galM, glk, pgm, pgi, pfkA, pfkB, fbp, fbaB, gapC, gpmA, gpmB, pykA, pykF, ppsA, zwf, pgl, gnd, rpe, rpiA, rpiB, tktA, tktB, talA,* and *talB*). We compared the predicted flux changes using ΔFBA with the differences of 46 measured metabolic fluxes along the central carbon metabolism by incorporating the enzyme expression obtained from RT-PCR. Uncentered Pearson correlations, NRMSE, and sign accuracy of the flux change predictions are depicted in **Figure 1B** and **C**, indicating a general agreement between the predicted flux changes and the differences in the measured fluxes. We compared the accuracy of flux change predictions by ΔFBA and REMI ^30^. As illustrated in **Figure 1B** and **C**, ΔFBA generally outperforms REMI in predicting the flux changes by having lower NRMSE, higher Pearson correlations, and higher sign accuracy. The results using the whole-genome gene expression profiles for a subset of perturbation experiments are comparable with those using RT-PCR data above (see also **Supplementary Figure S2** and **Supplementary Figure S3**).

**Figure 1:**
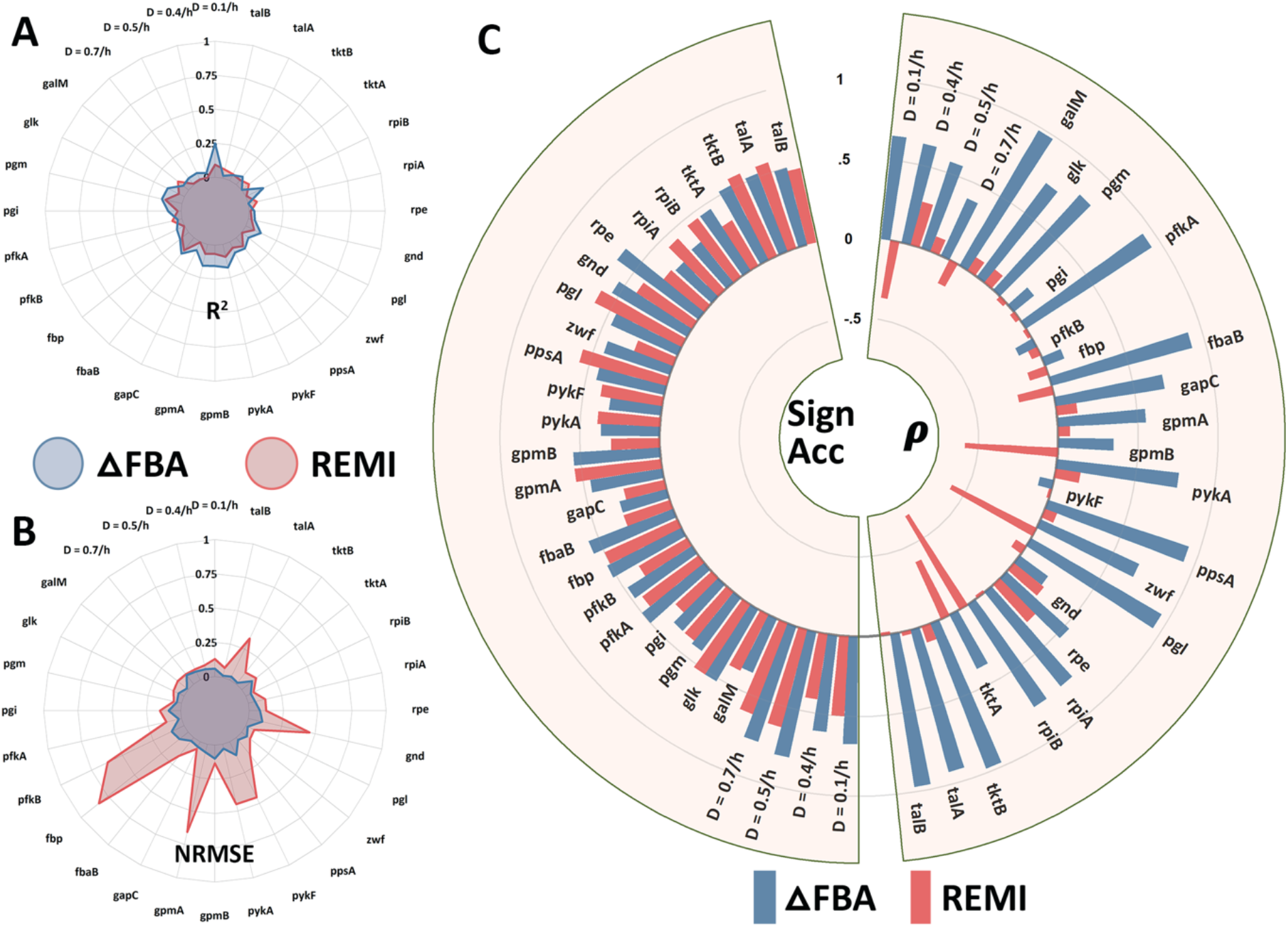
Comparison of the performance of ΔFBA and REMI in predicting *E. coli* metabolic response to environmental (dilution rates) and genetic (single gene deletions) perturbations. (A) The coefficient of determination (R^2^) between the measured flux ratios and the reaction expression ratios shows that differential gene expressions do not directly inform the metabolic flux changes. (B) Normalized Root Mean Square Error (NRMSE) of the predicted flux change – smaller indicates higher accuracy (mean NRMSE: ΔFBA = 0.14, REMI = 0.54). (C) Directional (Sign Accuracy) agreement and uncentered Pearson’s Correlation Coefficient (ρ) between the predicted and measured flux difference for the 46 reactions (mean sign accuracy: ΔFBA = 0.49, REMI = 0.43; mean ρ: ΔFBA = 0.61; REMI = −0.06).

Another study, carried out by Gerosa *et al.* ^35^, looked at how *E. coli*’s central carbon metabolism adapts to 8 different carbon sources: acetate, fructose, galactose, glucose, glycerol, gluconate, pyruvate and succinate. The study generated ^13^C metabolic flux, metabolite concentration and microarray gene expression data from exponentially growing *E. coli* under each carbon source. The study found that only a small subset of the numerous transcriptome changes translates to notable shifts in the corresponding metabolic fluxes, indicating non-trivial relationships between transcriptional regulations and metabolic fluxes. We applied ΔFBA to predict flux changes between every pair of the carbon sources, treating one as the perturbation and another as the control condition. **Figure 2** describes the good agreement between the flux change predictions by ΔFBA with the measured differences of 34 metabolic fluxes between any two carbon sources, specifically in terms of uncentered Pearson correlation (mean: 0.61), NRMSE (mean: 0.15), and sign accuracy (mean: 0.66). The findings from Ishii *et al*. and Gerosa *et al*. highlight the ability of ΔFBA in accurately predicting metabolic flux alterations using transcriptomic data for both environmental (e.g., dilution rates, carbon sources) and genetic perturbations.

**Figure 2:**
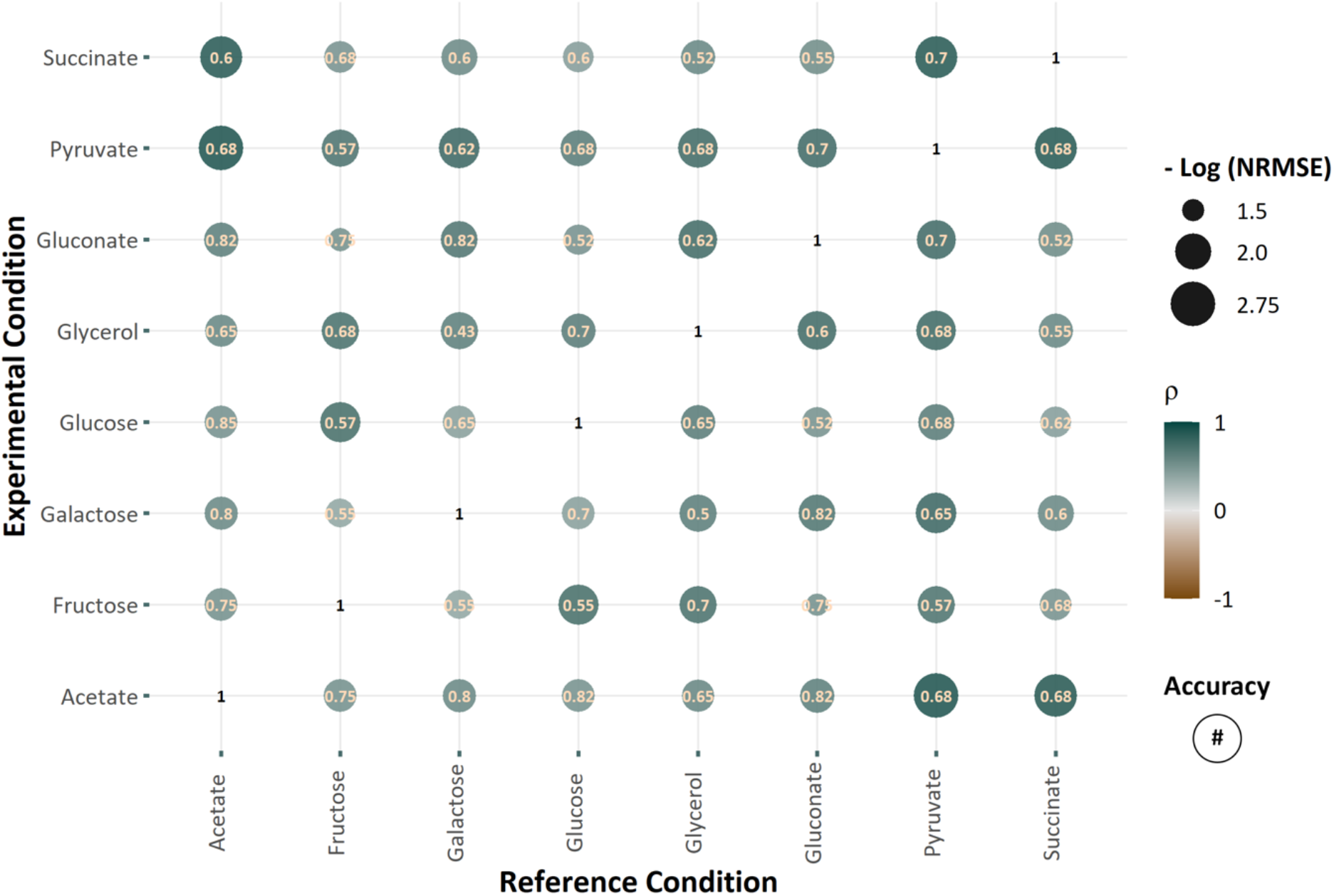
Prediction of metabolic flux changes in *E. coli* caused by changes in the carbon source using ΔFBA. The horizontal axis reports the reference carbon source (control) and the vertical axis shows the altered (perturbed) carbon source. Uncentered Pearson’s Correlation Coefficient (*ρ*) is shown by the color of the markers. NRMSE is represented by the size of the markers – the larger the markers, the smaller is the NRMSE. Finally, the directional (sign) accuracy of the flux perturbation predictions is shown by the numbers inside the markers.

### Dysregulation of Skeletal Muscle Metabolism in Type – 2 – Diabetes

In this case study, we looked at metabolic alterations of human muscle using the myocyte GEM *iMyocyte2419* ^36^ and gene expression datasets from two type-2 diabetes (T2D) studies, one by van Tienen *et al*. ^41^ and another by Jin *et al*. ^42^. In long term T2D patients compared to age-matched cohort, van Tienen *et al*. ^41^ reported the downregulation of gene expression related to substrate transport into mitochondria, conversion of pyruvate into acetyl-CoA, aspartate-malate shuttle in mitochondria, glycolysis, TCA cycle, and electron transport chain. Similarly, Jin *et al*. ^42^ reported a significant enrichment of pathways involved the oxidative phosphorylation among the downregulated genes in their T2D cohort compared to control. Jin *et al*. ^42^ further identified the transcription factor SRF and its cofactor MKL1 among the top-ranking enriched gene sets with increased expression. But, the correlation between the differential gene expressions in the two studies is only modest. ^36^

We applied ΔFBA to predict the flux changes based on the differential gene expressions in each of the two studies above (see Methods). We grouped the reactions based on whether the predicted flux changes are positive or negative, denoted by up- and down-reactions, respectively. We performed metabolic subsystem enrichment analysis using the subsystems defined in myocyte specific GEM *iMyocyte2419* ^36^ to identify over-represented metabolic subsystems among the up- and down-reactions (see Methods). As summarized in **Figure 3**, the enrichment analysis of metabolic changes in the van Tienen *et al*. study shows a significant over-representation of ß-oxidation and BCAA (branched-chain amino acids) metabolism among the down-reactions, and of extracellular transport and lipid metabolism among the up-reactions. The enrichment analysis of flux changes in the Jin *et al*. study also indicates an over-representation of lipid metabolism among the up-reactions in T2D patients, as well as an over-representation of ß-oxidation pathway among the down-reactions (see **Figure 3**).

**Figure 3:**
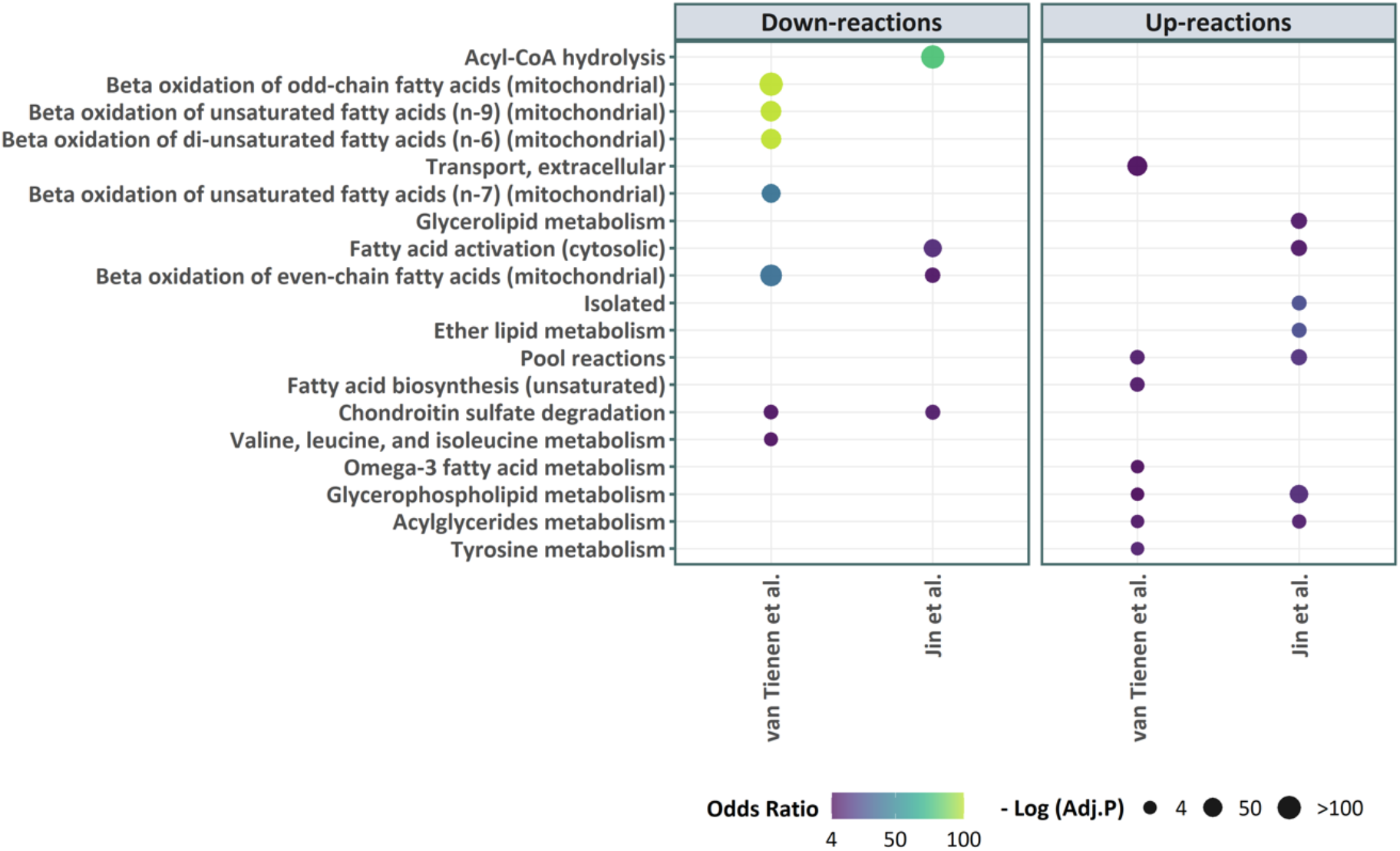
Enriched metabolic subsystems (FDR<0.05) among the in T2D patients based on flux changes predicted using ΔFBA. The flux changes were computed based on the transcriptome datasets from two T2D studies: van Tienen *et al*. ^41^ (GSE19420) and Jin *et al*. ^42^ (GSE25462). The statistical significance of the over-representation is shown by the size of the markers – larger markers have smaller adjusted *p*-values – while the odds ratio is shown by the color of the markers.

Furthermore, we evaluated the change in the flux throughput for every metabolite irrespective of its compartmental location, more specifically by computing the change in the total production flux of each metabolite. Metabolites with a large change in the flux throughput are of particular interest for disease biomarkers. In the following, we focus on metabolites that have a flux throughput change above a threshold (**|**Δ***ν*| > *ε***; ***ε* = 1**), and capture intermediary metabolites that participate in linear reaction sequences. **Figure 4** shows the flux throughput changes predicted by ΔFBA for various metabolites. Among the metabolites with a large drop in the flux throughput in both studies are Coenzyme A (CoA), Acetyl-CoA and AMP (Adenosine monophosphate), all of which have been previously identified as metabolite reporters of diabetes ^36,45^. Other metabolic biomarkers that have been previously proposed for T2D, such as repression of FAD (Flavin adenine dinucleotide), FADH^2^ and NADH by van Tienen *et al*. study ^41^ and increased glycerol by Jin *et al*. study ^42^, are confirmed by ΔFBA (see **Figure 4**). Väremo *et al.* ^36^ had also identified the metabolites above as T2D reporters via meta-analysis of numerous T2D datasets—including the two studies used here—and topology-based analysis using the GEM *iMyocyte2419.* Besides these confirmatory observations, our ΔFBA results further suggest that arachidonate and palmitate are candidate metabolic reporters for T2D, both of which have a significant positive flux throughput change in the two T2D studies (see **Figure 4**). The roles of arachidonate and palmitate in the progression and cause of T2D have also been previously investigated ^46–48^. The results above showcase the ability of ΔFBA in elucidating metabolic flux alterations in a complex human GEM and identifying key metabolites of interest in human diseases.

**Figure 4:**
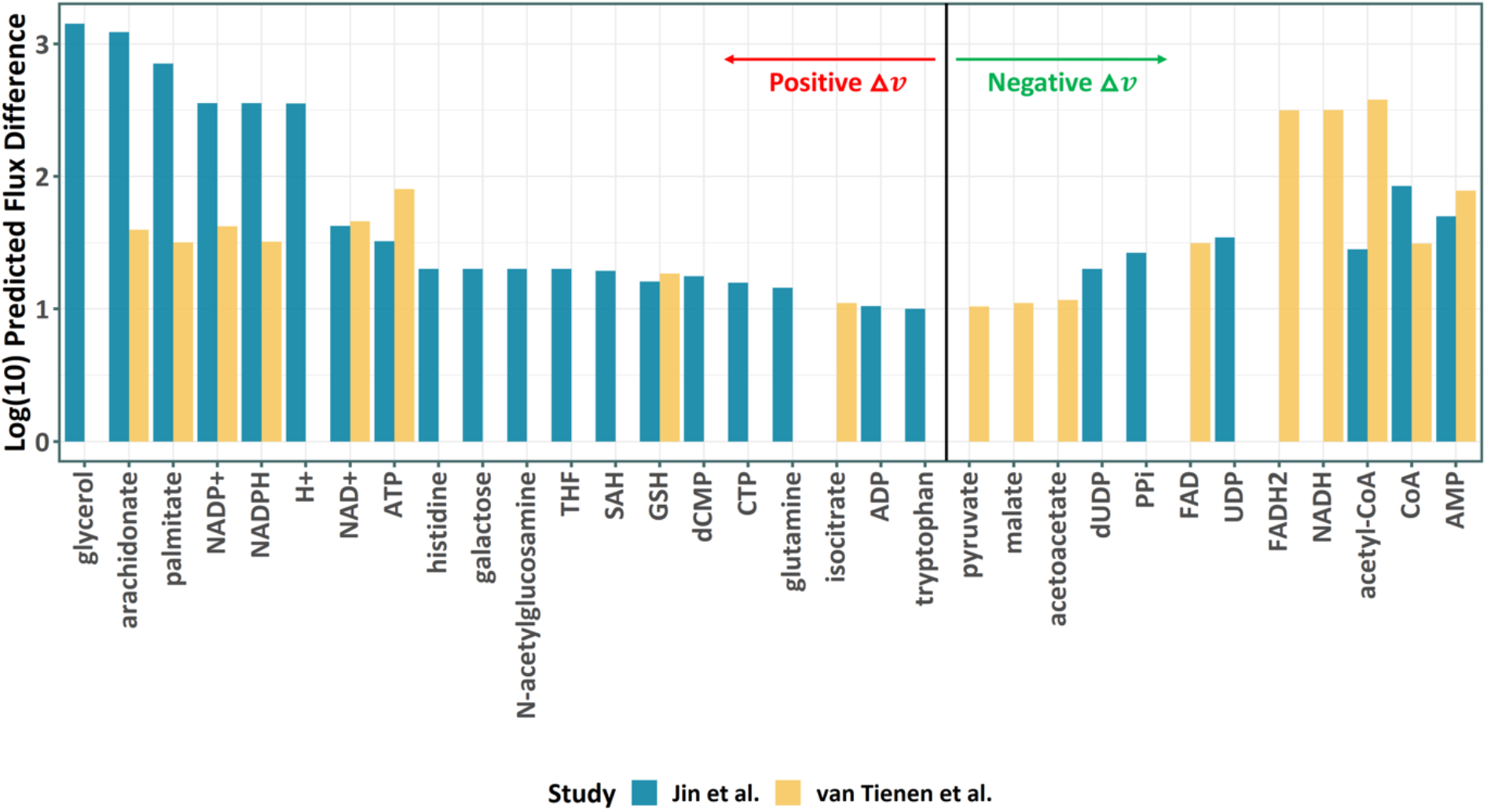
Alterations in metabolite flux throughput in T2D patients as predicted by ΔFBA.

## Discussions

GEMs and constraint-based modeling using FBA and the myriad FBA variants have proven to be important enabling tools for establishing genotype-phenotype relationship ^10,49,50^. The increasing availability of omics data have driving the development of FBA-based strategies that are able to use such data to improve the accuracy of predictions of intracellular metabolic fluxes. In this work, we present a new FBA-based method, called ΔFBA, built for the purpose of analyzing the metabolic alterations between two conditions given data on differential gene expression. ΔFBA does not require the specification of the metabolic objective, and thus, eliminates any potential pitfalls that are associated with an incorrect selection of this objective. Note that ΔFBA does not generate the flux prediction for a given condition; rather, the method produces differences of metabolic fluxes between two conditions. Differential flux predictions are indispensable in formulating hypothesis and in understanding the physiological response of cells to changes in the environment. ΔFBA can be easily integrated and have been tested to work with the widely popular COBRA toolbox.

We showed the applicability and performance of ΔFBA for predicting metabolic flux changes in an array of experimental perturbations and in both simple prokaryotic *E. coli* and complex multicellular human muscle cells. In comparison to a related method REMI ^30^, ΔFBA show a markedly better accuracy in prediction the magnitude and direction of metabolic flux changes in *E. coli*. Further, the application of ΔFBA to two T2D studies shed light on the rewiring of muscle metabolism associated with type-2 diabetes that leads to the repression of ß-oxidation and activation of glycerolphospholipids, pointing to increased lipid metabolism in the T2D patients. Interestingly, serum metabolic profiling of T2D patients showed increased glycerophospholipids when compared to healthy controls ^51^. Besides, clinical and experimental studies have demonstrated the association between phospholipids and insulin resistance ^52^. Furthermore, by looking at the changes in the flux throughput of metabolites, the results of ΔFBA suggest two fatty acids, arachidonate and palmitate, for candidate biomarkers of T2D.

There are several limitations of ΔFBA, the most obvious of which is that the method does not produce flux predictions for the conditions under comparison (perturbed vs. control). If the values of the metabolic fluxes are desired, one can use a FBA-based method, for example parsimonious FBA ^53^, to evaluate metabolic fluxes for one of the conditions (perturbed or control) – preferably one that is more well characterized (e.g., more experimental data, more obvious metabolic objective) – and compute the metabolic fluxes for the other condition by combining this flux prediction and the flux changes from ΔFBA. Also, in the formulation and the application of ΔFBA in this work, we considered only differential gene expression data. But the method can also accommodate other omics dataset, such as proteomics, by appropriate mapping of the data to changes in reaction expressions. Metabolomics data can also be accommodated in ΔFBA via thermodynamics constraints, as done in REMI ^30^, in which certain reactions can only proceed in one direction.

## Conclusion

In this work, we address the challenge of studying metabolic flux alterations in organisms as a result of genetic or environmental changes. Our versatile method ΔFBA provides a set of functions for evaluating metabolic flux differences between two conditions using genome-scale metabolic models and differential gene expression data. The computational tool eliminates the need for assuming a metabolic objective by exploiting the fact that the flux differences also satisfy the same flux balance equation used in standard FBA. As demonstrated in several case studies, ΔFBA provides accurate and biologically relevant predictions of metabolic alterations caused by environmental, genetic, and disease-related perturbations. With increasing research efforts directed toward the integration of omics data with biochemical network models, tools such as ΔFBA represents an important advancement in this direction. The MATLAB implementation of ΔFBA is freely available on https://github.com/CABSEL/DeltaFBA.

## Acknowledgments

The authors would like to acknowledge the Swiss National Science Foundation for funding support (grant number: 163390) and the University at Buffalo’s Center for Computational Research for computational support.

