## Supplementary Materials for "ΔFBA: Predicting metabolic flux alterations using genome-scale metabolic models and differential transcriptomic data"

Alternative Threshold Criteria of Minimum Flux Change Magnitudes

The thresholds for the minimum magnitude of positive and negative flux changes, $\mu_i$ and $\eta_i$, respectively, are user-defined parameters which can be a constant value $\varepsilon$, or a value that scales with the fold-change reaction expression $e_i^{P/C}$. If

$$\mu_i = \varepsilon \quad (1)$$

$$\eta_i = \varepsilon \quad (2)$$

When $z_i^U = 1$, $\Delta v_i$ should only be greater than $\varepsilon$ (default = 0.1) and when $z_i^D = 1$, $\Delta v_i$ will be less than $-\varepsilon$. However

$$\mu_i = \varepsilon e_i^{P/C} \quad (3)$$

when $z_i^U = 1$, $\Delta v_i$ will be scaled proportional to the upregulated reaction fold change. Similarly, by setting

$$\eta_i = \varepsilon / e_i^{P/C} \quad (4)$$

when $z_i^D = 1$, $\Delta v_i$ takes a negative value proportional to the downregulated reaction. Such scaling introduces a more stringent constraint on $\Delta v_i$.

We tested the threshold prescribed by Equation (3)-(4) above for the Ishii et al. study, and compared the predicted flux changes $\Delta \boldsymbol{v}$ with those obtained with a constant threshold in Equations (1)-(2). The introduction of thresholds that scales proportionally with the fold changes of the reaction expression did not significantly alter the predictions of $\Delta \boldsymbol{v}$, as shown in **Figure S1**. The difference in the sign accuracy and the correlation were only marginal (Mean correlation coefficient (ρ): Eqs. (1)-(2) threshold = 0.61, Eq. (3)-(4) threshold = 0.58; Mean directional accuracy: Eqs. (1)-(2) threshold = 0.49, Eq. (3)-(4) threshold = 0.48; Mean NRMSE: Eqs. (1)-(2) threshold = 0.14, Eq. (3)-(4) threshold = 0.16).

Supplementary Figure S1

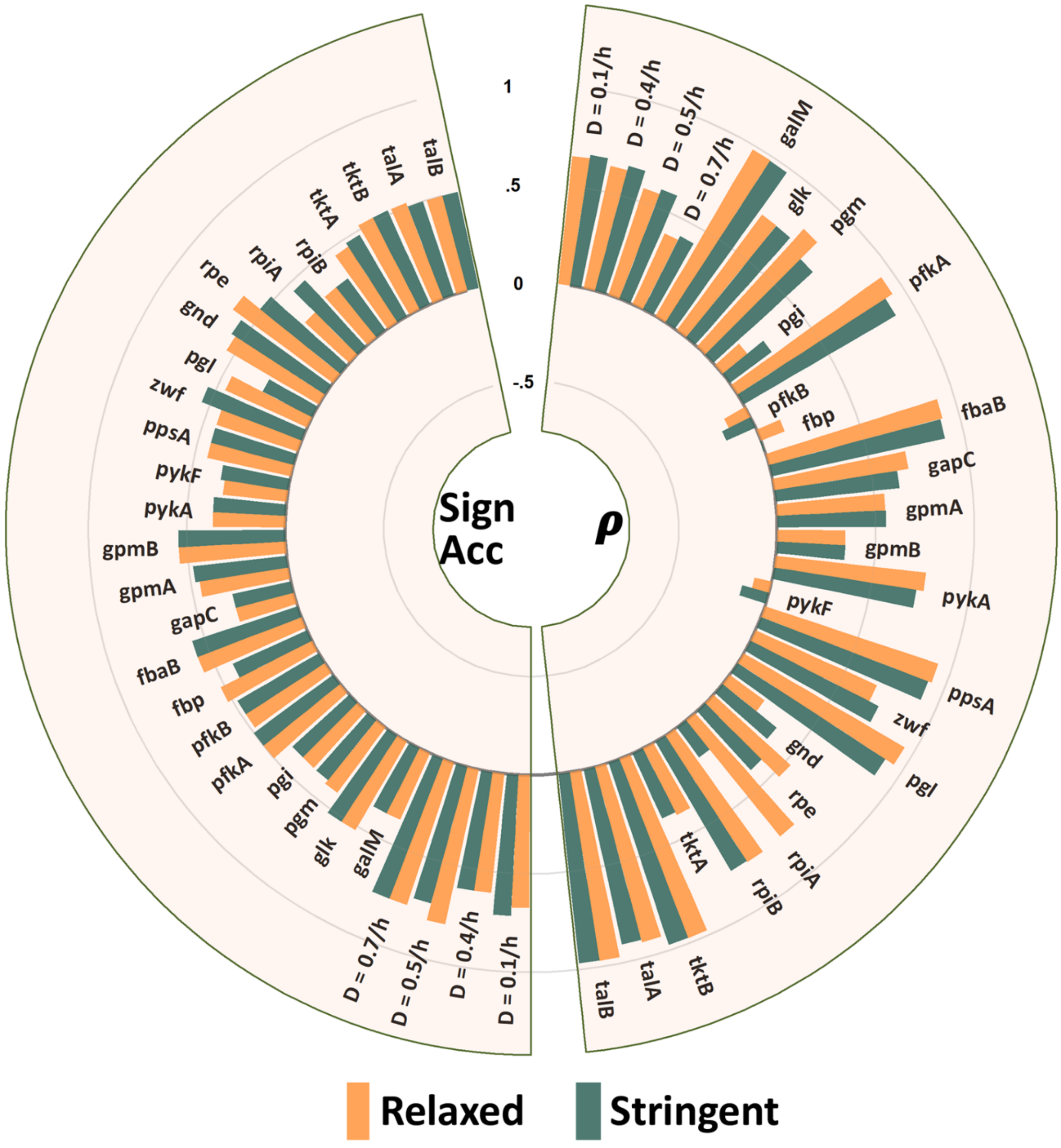


Figure S1: Comparison of ΔFBA predictions of *E. coli* metabolic response to environmental and genetic variations using constant and fold-change criteria for minimum flux change magnitude. The accuracy of the flux predictions is assessed by calculating the Uncentered Pearson's Correlation Coefficient (ρ) and the directional (sign) agreement between the predicted flux difference and the measured flux change for 46 reactions. The difference in flux change predictions between the two thresholds is insignificant.

Supplementary Figure S2

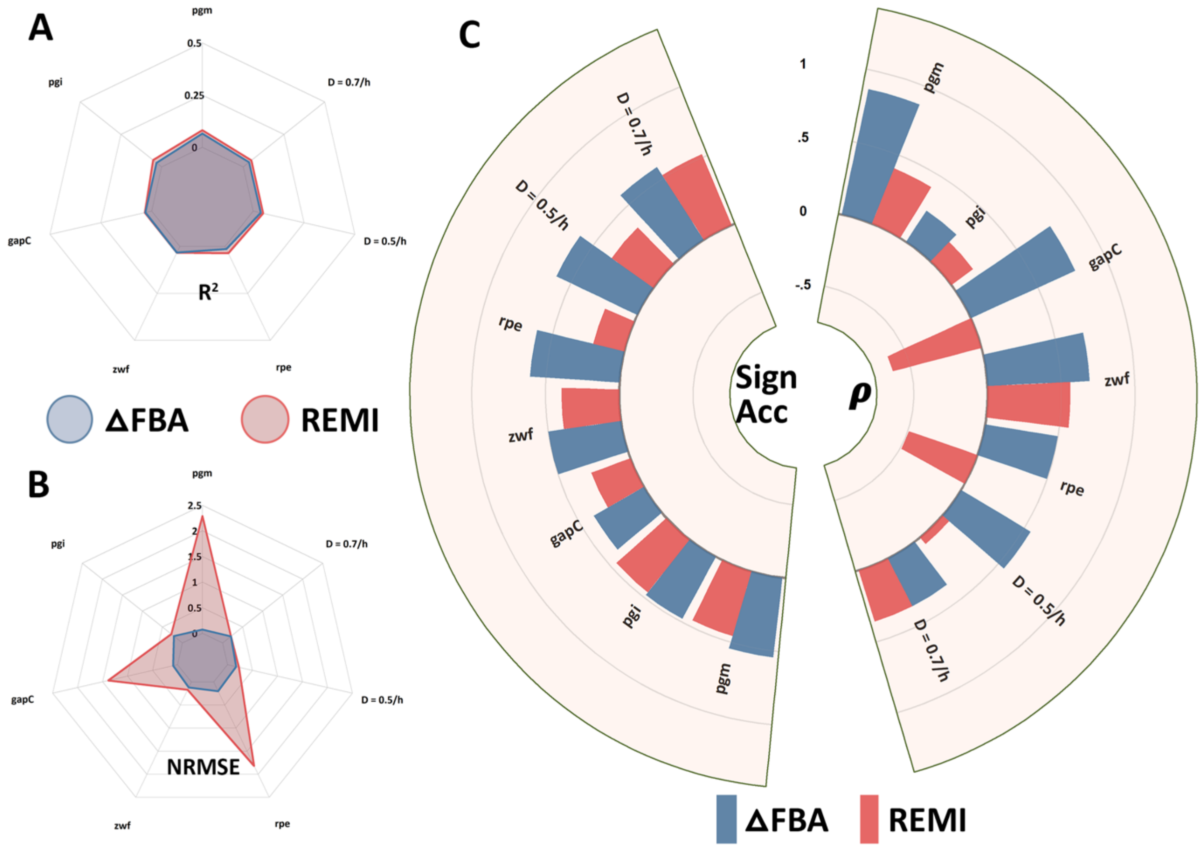


Figure S2: Comparison of the performance of ΔFBA and REMI in predicting *E. coli* metabolic response to 2 dilution rate and 5 single-gene deletion perturbations using whole-genome transcriptome data. (A) The coefficient of determination ($R^2$) of the measured flux ratios and the reaction expression ratios remains close to zero. (B) Normalized Root Mean Square Error (NRMSE) of the flux difference normalized over the measured flux difference (Mean NRMSE: ΔFBA = 0.15; REMI = 0.91). (C) Directional (Sign Accuracy) agreement and uncentered Pearson's Correlation Coefficient (ρ) between the predicted and measured flux difference for the 46 reactions (mean sign accuracy: ΔFBA = 0.53, REMI = 0.38; mean ρ: ΔFBA = 0.57; REMI = 0.06).

Supplementary Figure S3

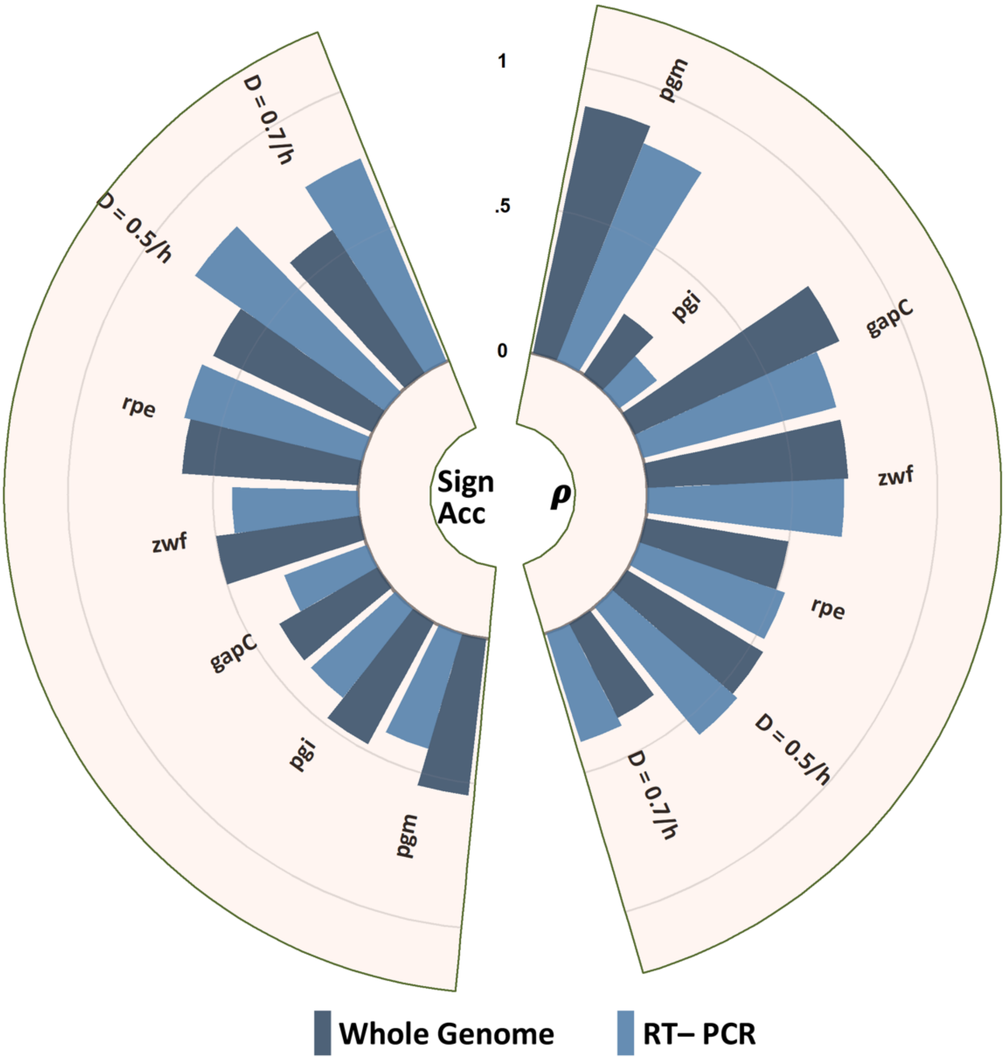


Figure S3: Comparison of ΔFBA performance in predicting *E. coli* metabolic flux changes between using whole-genome transcriptome data and using RT-PCR mRNA data. Directional (Sign Accuracy) agreement and uncentered Pearson's Correlation Coefficient (ρ) between the predicted and measured flux difference for the 46 reactions show little differences between the two types of transcriptomic data (mean sign accuracy: whole-genome = 0.53, RT-PCR = 0.53; mean ρ: whole-genome = 0.57; RT-PCR = 0.54).
